# Reactive Initiation Training is more effective than Shuttle Run to improve the on-court agility of novice badminton players

**DOI:** 10.1101/331058

**Authors:** Minkai Dong, Jidong Lyu, Thomas Hart, Qin Zhu

## Abstract

**Purpose:** Despite its well-known importance in sports, agility is ambiguously defined and lack of research. Shuttle Run (SR) is commonly used to improve the on-court agility of badminton players. Reactive Initiation Training (RIT) contrasts SR in that it only demands rapid generation of initiation step toward the direction of shuttlecock. The current study compared SR with RIT to determine which one is more effective for improving on-court agility of novice badminton players.

**Method:** 20 novice badminton players were split in half to receive either RIT or SR on court for five days. Before and after training, participants were assessed on their ability to intercept the shuttlecocks randomly thrown by a coach to six corners of the court with and without visual occlusion of the coach. All trials of interception were recorded for video analysis of initiation time, running time and total time.

**Results:** The mean total times were greater with visual occlusion and varied systematically with the position of interception. Both training methods shortened the mean running time, however, only RIT additionally reduced the initiation time and its proportion on those time-consuming positions in the occluded condition.

**Conclusion:** RIT is more effective than SR to improve the on-court agility of novice badminton players.

## Introduction

Agility has been traditionally referred to as motor quickness including the ability to generate explosive power^1,2^ and the ability to change direction rapidly^3,4^. However, a comprehensive definition of agility suggests that the cognitive quickness related to perceptual and decision-making skills is another key element constituting agility^5,6^, and should be considered in developing sport-specific agility test and training^7,8^. In invasion sports where opponents attempt to invade each other’s territory to gain advantages, the cognitive element of agility test has been shown sensitive to discriminate between high-level and low-level athletes^9^. While the plyometric training seems to be effective to improve the change-of-direction speed, the one-on-one training or small-sided games appear to be beneficial for improving the reactive agility^10^.

Badminton is a net/wall game with a net diving players’ territory. While players constantly use directional shots to outscore opponents, they must also return the opponent’s shots by running rapidly and repeatedly on court with change of direction to intercept the shuttlecock. Given the short shuttlecock flight time in a rally^11,12^, the player typically has less than one second to react and run to complete the interception. Therefore, badminton demands on-court agility that includes both cognitive and motor quickness and having the ability to anticipate the shot will greatly ease the challenge to improve the agility^13^.

Various methods have been implemented to train on-court agility in badminton^14,15,16,17^. Shuttle run (SR) has been a popular and badminton-specific agility training that involves constant change of speed and direction to reach the corners of the court in a predetermined order. As a high-intensity interval training, SR is preferred by researchers and coaches because it resembles the characteristics of badminton including the actions during play, temporal structure of playing and rest time, and the resultant heart rate and lactate concentration after play^11,18^. However, SR has a limited transfer to the real game due to the missing reactive component and randomness of movement pattern during training. Although a recent study included those missing factors in SR^19^, only the day-to-day variation of performance was evaluated, and a valid on-court agility test was warranted to determine the effectiveness of SR training.

Despite the use of video-based judgment tasks to train badminton players’ perception and anticipation^20,21^, the effect of such a training on the on-court agility remains undetermined due to its lack of demand for on-court actions. It has been suggested that the sport-specific reactive agility training should demand high information-movement coupling and replicate the game situations^22^. Therefore, the reactive initiation training (RIT) that demands rapid generation of the initiative step in response to the perceived direction of shuttlecock could serve to improve the reactive agility of badminton. Compared to SR, RIT challenges the cognitive processing more than the physical effort, therefore is less intensive.

Using an on-court agility test simulating the game play in badminton, the current study is aimed to compare SR with RIT (both implemented on court) to determine which one is more effective to improve the on-court agility of novice badminton players. Since the on-court agility test demands both cognitive and motor quickness and each training is specific to improve one of them, the contribution of each training to the general agility and the potential transfer of training can both be evaluated in this study.

## Methods

Based on a pilot study that achieved an intermediate effect size for the interested factors in a mixed design ANOVA, 20 novice badminton players (split in gender) were recruited for the current study. They were all college students enrolled in the 2016 Fall class of *Introduction to Badminton* at Shanghai University of Finance & Economics. All participants were right-handed and judged by the instructor to have limited experience and playing skills in badminton. Written informed consent was obtained from each participant before their participation in the study. The experimental protocols were approved by the IRB committees of both Shanghai University of Finance & Economics and University of Wyoming.

All participants were assessed with an on-court agility test before and after receiving the agility training. The on-court agility test involved intercepting a randomly thrown shuttlecock on a regular badminton court as quickly and accurately as possible. The center line was marked by a small piece of blue tape 1.95m from the short service line, and the participant was asked to occupy the marked position with a racquet held in hand. A certified badminton coach stood behind the net on the other side of the court, throwing a shuttlecock from the center of the net. The shuttlecock was thrown randomly to six positions representing the six corners of the court, and they were labeled counterclockwise 1 to 6 as seen in Fig 1.

**Fig 1.**
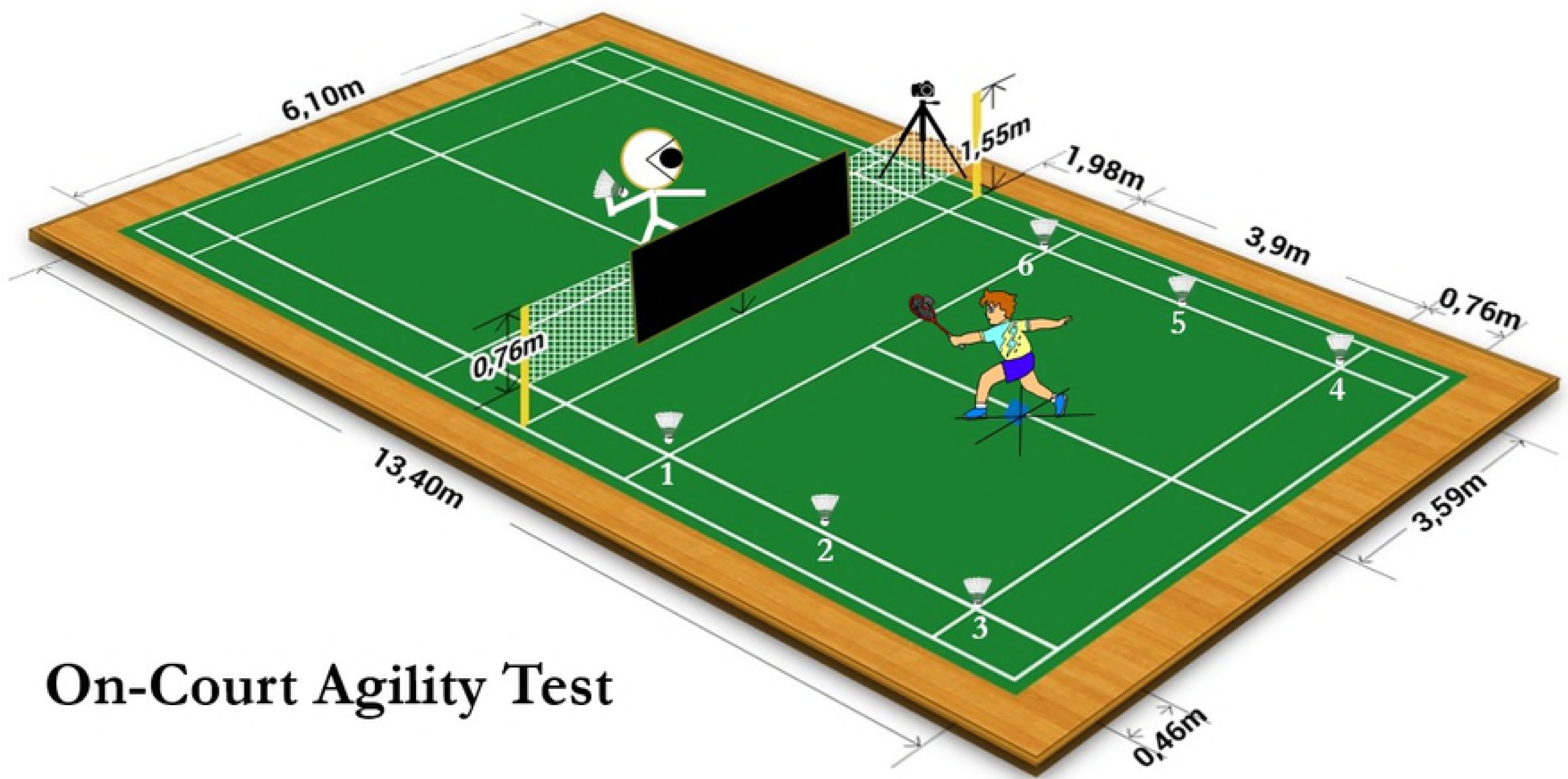
The diagram depicting the on-court agility test. The coach was given a random order sheet to throw the shuttlecock for 18 times so that each of the six corners were randomly attempted by the player three times. In each trial, the participant was instructed to react by running to intercept the shuttlecock using the racquet. Jumping or diving to intercept was not allowed and successful return of the shuttlecock over the net was encouraged but not required. To limit the factor of technique, any part of the racquet could be used for interception, and in case the shuttlecock was not intercepted before landing, the participant had to tap the landed shuttlecock using the racquet as soon as possible to complete the interception. The participant was given ample recovery time to return to the starting position before the next trial. The on-court agility was tested twice with and without visual occlusion of the coach. In the occluded condition, a black plastic curtain was hung over the net to mask the vision of the entire opponent’s court. Therefore, the coach and his throwing motion were unseen by the participant, and the participant had to respond upon the first sight of the thrown shuttlecock. The order of receiving visual occlusion or not was counterbalanced among participants. A GoPro Hero 5 camera was used to record the participant’s performance in the agility test. The camera was set with a recording rate of 120fps in resolution of 1080p before being fixated on a tripod and positioned next to a net pole facing the participant. The camera view captured the entire side of the participant’s court as well as the net such that the release of the shuttlecock from the coach’s hand, the movement of the participant on court, and the interception of the shuttlecock could all be observed.

Following the pre-test of on-court agility, participants were randomly divided into two groups to receive a prescribed agility training with four bouts each day for five days. The gender was balanced so that there were five males and five females in each group. The SR group focused on improving the running speed to reach different corners of the court. The four bouts of SR in each day were separated by a five-minute interval to allow for full recovery from maximum effort work. At each corner of the court, four up-turned shuttlecocks were aligned to be perpendicular to the line connecting the corner and the marked starting position. Each bout of running was timed for 30 seconds. The participant should run repeatedly from the center starting position to knock down the shuttlecocks using the racquet hand in a predetermined order (position 6, 1, 5, 2, 4, and 3). The total number of knockdowns and the time taken to knock down the 12^th^ shuttlecock in each bout were recorded, and the participants were encouraged to increase the number and shortened the time with each bout.

The RIT group focused on improving the reactive initiation toward different corners of the court. The day training consisted of four bouts of reaction to the screen-displayed arrows with two minutes break between bouts. In each bout, the participant stood at the starting position with bent knees to prepare for foot initiation. A large pullup projector screen (84inch diagonal) was set up in the center of net area facing the participant, and a projector was connected to a laptop computer to project the stimulus onto the screen. In each trial, participants first attended to a white dot displayed in the center of screen in black backdrop representing the starting position, shortly after, a white arrow appeared to point at one of six corners from the white dot. Upon seeing the arrow, participants should initiate a step with the racquet foot toward the indicated corner as soon as possible. The initiation step toward the left-back corner could be taken by turning the body clockwise or counterclockwise, whichever was easier for the participant. The 30 trials of arrow presentation were programmed by E-Prime software (Ver 3.0), where the directions of the arrows and the fore-period of the presentation were randomized. The interval between trials was fixed to be 5 seconds allowing sufficient time for the participant to recover to the starting position. The day training was recorded by a GoPro camera and the times taken to complete the initiation step was obtained through tracking the converted videos in MaxTRAQ software (Ver 2.8.4.3). Participants were provided this information to improve in the next day training.

Following the five days of training, all participants were examined with the same on-court agility test. All the recorded videoclips were converted to maintain the original resolution and recording rate for tracking in MaxTRAQ to identify three critical events: A) the release of the shuttlecock from the coach’s hand, B) the initiation of the racquet foot toward the correct direction, and C) the completion of the interception. Subsequently, the initiation time was calculated as the interval between event A and B, the running time as the interval between event B and C, and the total time as the interval between event A and C. Therefore, the proportion of initiation time was the ratio between initiation time and total time. All the dependent variables were submitted to the 2 (group) x 2 (phase) x 2 (occlusion) x 6 (position) mixed design ANOVA to examine the effects of between-subject variable (group), within-subject variables (phase, occlusion and position), and their interaction, followed by the necessary post-hoc analyses. The statistical significance was set at p < 0.05.

## Results

As seen in Fig 2, the mean total times varied with positions in a “W” shape, and significantly reduced after training for all participants. Participants were generally slower with visual occlusion.

**Fig 2.**
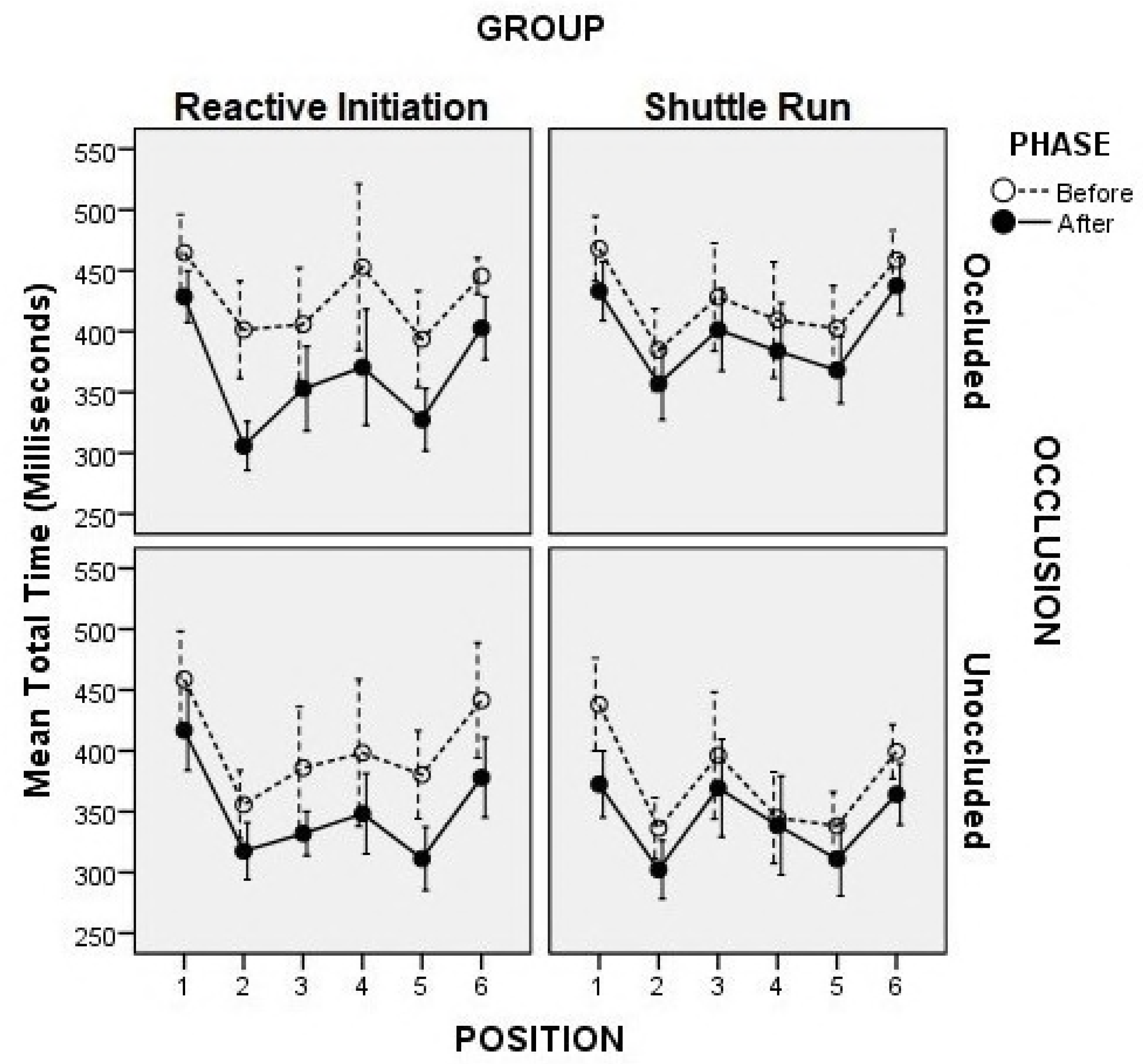
The mean total time as a function of training group, training phase, position and occlusion. The ANOVA on the total times showed significant main effects for phase (F_1,18_ = 39.55, p < 0.001), occlusion (F_1,18_ = 56.95, p < 0.001), and position (F_15,90_ = 16.05, p < 0.001). The Tukey’s post-hoc analysis revealed that the mean total times on position 1 and 6 were significantly longer than that on position 3 and 4 (p < 0.05), and even longer than that on position 2 and 5 (p < 0.05). The interaction between group and occlusion was also significant (F_1,18_ = 12.31, p < 0.01). As revealed by the simple main effect analysis, the occlusion effect was significant for the SR group (F_1,234_ = 27.19, p < 0.001), but marginal for the RIT group (F_1,234_ = 3.63, p = 0.058).

As seen in Fig 3, the initiation times and its proportion varied with positions systematically and reduced after training only for the RIT group in the occluded condition.

**Fig 3.**
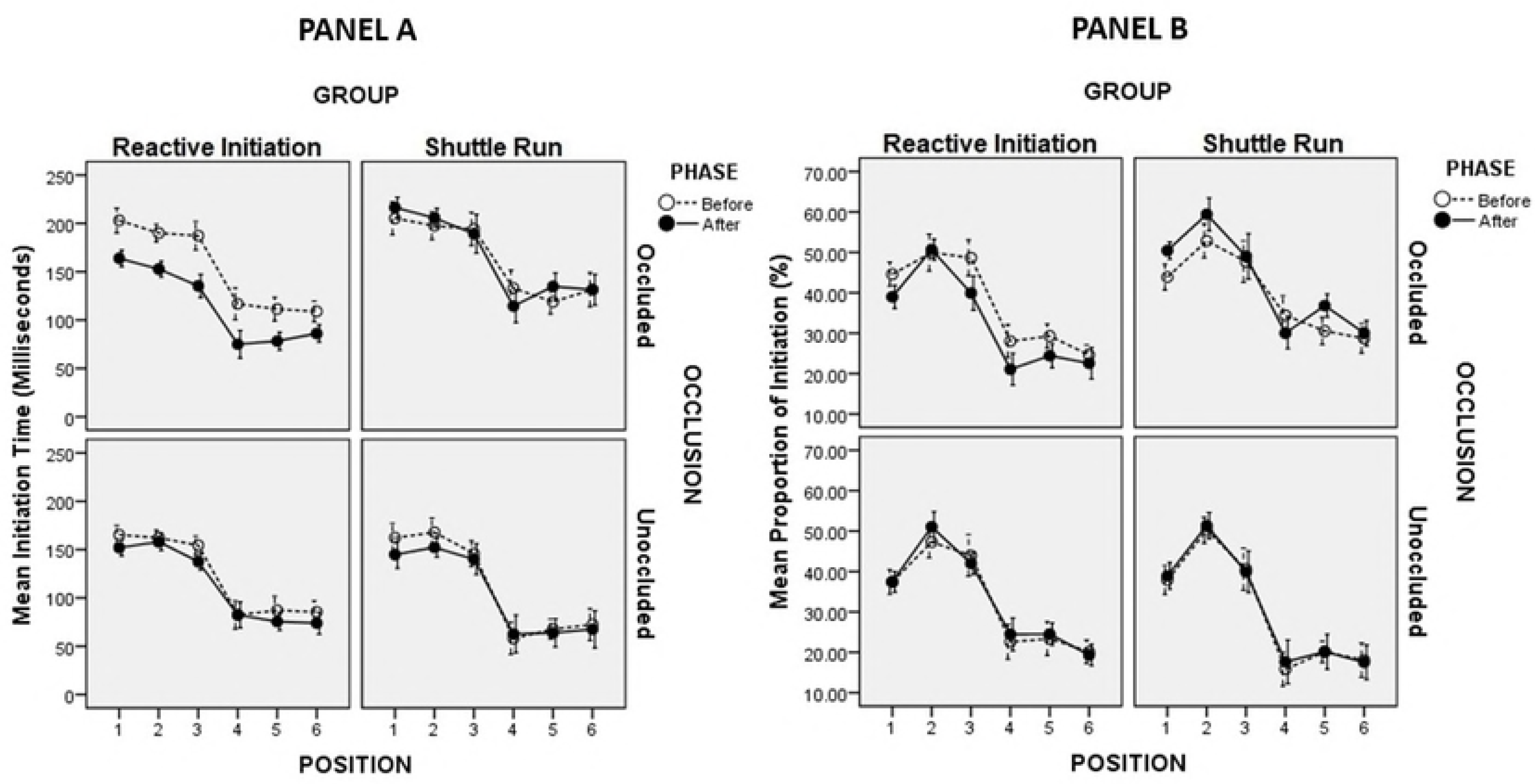
The mean initiation time and its proportion as a function of training group, training phase, position and occlusion. The ANOVA on the initiation times showed significant main effects for phase (F_1,18_ = 7.18, p < 0.05), occlusion (F_1,18_ = 287.75, p < 0.001), and position (F_5,90_ = 198.91, p < 0.001). The Tukey’s post-hoc analysis revealed that the mean initiation times on position 1, 2, 3 were significantly longer than that on position 4, 5, 6 (p < 0.05). Since the 4-way interaction was also significant (F_5,90_ = 2530.95, p < 0.01), a 2-way (phase x position) repeated measure ANOVA was performed separately for each group in each occlusion condition. The results showed that the phase (F_1,9_ = 30.24, p < 0.001), position (F_5,45_ = 83.82, p < 0.001) and their interaction (F_5,45_ = 2.76, p < 0.05) were all significant only for the RIT group in the occluded condition, while in other group-occlusion conditions, only the position effect was found significant (p < 0.001).

The ANOVA on the proportions of initiation time showed significant main effect for occlusion (F_1,18_ = 60.67, p < 0.001) and position (F_5,90_ = 118.01, p < 0.001), and a significant interaction among group, phase, and occlusion (F_1,18_ = 8.85, p < 0.01). The follow-up analyses revealed that the phase by position interaction was significant for both groups only in the occluded condition (F_5,45_ = 2.67, p < 0.05 for RIT group; F_5,45_ = 2.53, p < 0.05 for SR group). The simple main analyses further suggested that the RIT reduced the proportion of initiation time significantly on position 1, 3, 4 (p < 0.05), and marginally on position 5 (p = 0.059). In contrast, the SR increased the proportion of initiation time significantly on position 1, 2 (p < 0.05), and marginally on position 5 (p = 0.056).

The ANOVA on the running times only showed the significant main effect for phase (F_1,18_ = 16.65, p < 0.01) and position (F_5,90_ = 33.72, p < 0.001) without any interaction. The Tukey’s post-hoc analysis revealed that the mean running time were significantly greater (p < 0.05) on position 1, 4 and 6 than on position 3 and 5, and then on position 2.

## Discussion

Using the on-court training and testing paradigm, the current study examined the effectiveness of two training methods commonly used to improve the on-court agility of badminton with one focusing on the reactive initiation and the other on the change-of-direction speed. The results clearly suggested that novice badminton players benefitted more from the RIT than SR, even though both training helped to improve the overall performance on court.

The significant effect for position was found in all measures before and after training, suggesting that the on-court agility of badminton depends on the direction to take the shot. Overall, it takes longer time for the player to intercept the shuttle in the front and back corners as opposed to on the side. This makes perfect sense as clear and drop shots are often used by badminton players to move opponents around and slow them down, while smash is used to attack the side. However, the position effect differed for initiation time and running time. In general, players were faster initiating to the forehand side than to the backhand side, but took longer to intercept the shuttlecock on position 1, 4 and 6. In choice-reaction tasks, a faster reaction time is typically demonstrated when the stimulus and response are dimensionally compatible^23,24^. In this study, the initiation of the racquet hand and foot was spatially and conceptually mapped to the direction of thrown shuttlecock, therefore, the faster initiation on the forehand side is expected. However, after initiation, players took different times to intercept the shuttlecock. As revealed by the video analysis, players often initiated with a single lunge step to intercept the shuttlecock on position 2, 3 and 5 instead of taking multiple steps, consequently the greater running times on position 1, 4 and 6.

The occlusion effect was observed on total time, initiation time and its proportion, but not on running time, suggesting that players slowed down their initiation in response to the directional shots when the vision of the coach’s throwing motion was occluded. Studies have shown that people are adept at using the kinematic information in the pre-release throwing motion to predict the outcomes of throwing^25,26,27^. Therefore, the players in our study could anticipate the direction of the thrown shuttlecock and initiate faster with vision of the coach. When anticipation became impossible by visual occlusion, they slowed down the initiation but not the running speed.

The practice specificity effect was also evident. RIT was effective to reduce the initiation times on all positions in the occluded condition, while SR was effective to reduce the running times on all positions regardless of occlusion. Since the on-court agility of badminton demands the speed of both initiation and running, both RIT and SR should be recommended for agility training. However, the transfer effect was only observed for RIT. While the SR group did not improve the initiation at all, the RIT group improved the running speed along with the improved initiation. This is surprising because their training did not involve any running. The possible explanation is that RIT partially trained the muscles involved in running because the initiation required activation of the same muscles^28^, which yielded the neuromuscular adaptation to support ballistic movement following the initiation^29^. When the initiation times were normalized by the total times, it was evident that RIT specifically reduced the proportion of initiation time (leaving more time for running) on those time-consuming positions (position 1, 3, and 4), while SR increased the proportion of initiation time (leaving less time for running) on those time-saving positions (position 2 and 5).

## Conclusion

In sum, novice badminton players benefited more from RIT than SR by showing the reduction of both initiation time and running time after training. Therefore, RIT should be recommended as the main agility training for these players. Since SR was only effective to reduce the running time on court, it should be recommended as a supplementary agility training. However, considering that the significant reduction of initiation time following RIT was only observed in the occluded condition, the reported effect of RIT on the on-court agility is limited to the situation where the anticipation is impossible or to those players who have not yet developed adequate anticipation skills. Since anticipation is a critical component of reactive agility, incorporating the anticipation training in RIT (e.g. replacing the directional arrows with the opponent’s actions) should be promising to significantly enhance on-court agility in badminton.

## Practical Implications

- Reactive Initiation Training is a less intensive but effective agility training for novice badminton players
- Despite its popularity, Shuttle Run is an intensive but less effective agility training for novice badminton players
- Incorporating the anticipation training in Reactive Initiation Training is promising to significantly enhance on-court agility in badminton

## Acknowledgements

The authors want to thank all participants for their commitment of extra-curricular time and effort to complete the training and test.

## Supporting information

**Fig S1. The diagram depicting the on-court agility test.** Note: The blue taped spot in the center of the court marked the starting position in both test and training. During test, participants initiated from the starting position in each trial to intercept the thrown shuttlecock with and without visual occlusion. In Reactive Initiation Training, participants stood at the starting position to practice the initiation step in response to the randomly displayed directional arrows on a large pullup screen. In Shuttle Run, participants initiated from the starting position to knock down the up-turned shuttlecocks aligned at each corner in sequence of 6, 1, 5, 2, 4, 3 repeatedly.

**Fig S2. The mean total time as a function of training group, training phase, position and occlusion.** The empty dots connected by the dash line represent the mean total time on each position before training, and the filled dots connected by the solid line represent the mean total time on each position after training. Position 1 through 6 are respectively: the left-front corner, the left-side corner, the left-back corner, the right-back corner, the right-side corner, and the right-front corner, from the player’s perspective of view. The error bars represent the standard errors of the means.

**Fig S3. The mean initiation time and its proportion as a function of training group, training phase, position and occlusion.**The empty dots connected by the dash line represent the mean initiation time (in panel A) and its proportion (in panel B) on each position before training, and the filled dots connected by the solid line represent the mean initiation time (in panel A) and its proportion (in panel B) on each position after training. Position 1 through 6 are respectively: the left-front corner, the left-side corner, the left-back corner, the right-back corner, the right-side corner, and the right-front corner, from the player’s perspective of view. The error bars represent the standard errors of the means.

